# 3D Bioprinting Using Poly(ethylene-glycol)-dimethacrylate (PEGDMA) Composite

**DOI:** 10.1101/2023.10.19.562790

**Authors:** Shu-Yung Chang, Joseph Zhi Wei Lee, Anupama Sargur Ranganath, Terry Ching, Michinao Hashimoto

## Abstract

Recent progress in additive manufacturing has enabled rapid printing of bioinspired structures such as vasculature and alveoli using stereolithography (SLA) bioprinting. Bioinks for SLA often require synthetic polymers as additives to ensure the structural integrity of the printed cell-laden constructs. To this end, high molecular weight (MW) poly(ethylene-glycol)- diacrylate (PEGDA) (MW = 3400) is commonly used to enhance the mechanical property of crosslinked hydrogels, which requires in-house polymer synthesis or the acquisition of costly reagents. This research investigated the use of poly(ethylene-glycol)-dimethacrylate (PEGDMA) (MW = 1000) as a component of a composite bioink to enhance the mechanical properties of the SLA-printed constructs. We successfully demonstrated the fabrication of three-dimensional (3D) constructs with overhang and complex architecture, while human colorectal cancer cells (Caco-2) embedded in the crosslinked bioink exhibited the capability to proliferate on Day 6 of *in vitro* cell culture. Our study suggested PEGDMA as a viable alternative to high MW PEGDA used in SLA bioprinting. The accessibility to PEGDMA will facilitate the advance in 3D bioprinting to fabricate complex bioinspired structures and tissue surrogates for biomedical applications.

**Article Highlights:**

- Poly(ethylene-glycol)-dimethacrylate (PEGDMA) can be used in cell-laden bioprinting to enhance the mechanical property of bioinks.
- PEGDMA-based bioink was non-cytotoxic and conducive to cell proliferation.
- The facile preparation of PEGDMA composite ink will help to accelerate the research in tissue engineering via bioprinting.

## Introduction

This paper describes three-dimensional (3D) light-based bioprinting of large cell-laden tissue using a composite bioink consisting of fish gelatin methacryloyl (GelMA) and poly(ethylene-glycol)-dimethacrylate (PEGDMA). Ideal bioinks for SLA bioprinting should possess adequate mechanical properties to hold the printed structures while providing cytocompatible environments for cell growth and proliferation. In this work, we demonstrated that the GelMA-PEGDMA composite ink is suitable for printing 3D constructs with overhang and complex architecture using a digital light processing (DLP) printer. Crucially, we demonstrated that the GelMA-PEGDMA composite ink could be used for SLA bioprinting of cell-laden constructs with (1) sufficient mechanical strength tunable with the concentration of PEGDMA and (2) proliferative activities of the embedded cells. The developed bioink should pave an avenue for 3D bioprinting of large biomimetic tissues for applications in tissue engineering and organ-on-a-chip.

### Methods for bioprinting

3D bioprinting allows fabrication of complex and cell-laden biomimetic architecture. This ability gives it great potential in regenerative medicine, tissue engineering, and pharmaceutical sciences ^[1–2]^. Bioprinting can be broadly divided into two approaches: extrusion-based and light-based. In extrusion bioprinting, bioinks are loaded in a syringe and pneumatically extruded through a nozzle ^[3]^. Extrusion bioprinting has been shown to produce large-scale 3D biological constructs. However, the dispensing process of extrusion bioprinting inevitably exposes cells in the bioink to shear stress, which can be detrimental to the cells if the extrusion rate is not carefully selected ^[4–5]^. In addition, extrusion bioprinting involves point-by-point deposition of the bioink (usually as a continuous line), and the printing time increases with the size of the 3D construct. A prolonged printing time is not favorable for cell-laden bioinks; maintaining nutrients and oxygen supply to the cells during the printing process is crucial ^[6]^. As an alternative to extrusion-based methods, light-based methods, represented by SLA 3D printing, have been employed in bioprinting. SLA involves photopolymerization of resin in a reservoir in a layer-by-layer fashion using light projection. In SLA printing, thousands of pixels are independently controlled to project light of selected wavelength to polymerize 3D structures plane-by-plane. The difference in the printing mechanism allows SLA printing to produce large-scale 3D constructs faster than extrusion-based printing. In addition, high-resolution features can be readily achieved in SLA printing as its printing resolution is controlled by the size of each pixel in the digital light projection. SLA printing can be achieved using DLP, which is based on the processing of light through a digital micromirror device (DMD); this method is also called DLP 3D printing. In DLP printing using a commercial 3D printer, the pixel resolution can be as high as 27 µm ^[7]^. SLA printing, coupled with optimal concentrations of photoabsorbers, has been demonstrated to produce polyethylene glycol (PEG)-based mold to fabricate branching vascular constructs ^[8]^. Ultra-high precision of SLA at the nanometer range has also been demonstrated using technologies such as multiphoton lithography ^[9–10]^. Recent advances in SLA printing have enabled the fabrication of cells-laden constructs. For example, SLA bioprinting has been demonstrated to produce vascularized constructs with microvasculatures as small as 100 µm in channel diameter ^[11–12]^. To this end, it is crucial to employ suitable photocrosslinkable polymers and photoinitiators to ensure the viability and proliferation of cells in the 3D-printed constructs.

### Bioinks for SLA printing

Bioinks for SLA consist of cells, polymers, and photoinitiators. The polymers are photopolymerized by exposure to light during the printing, and the photopolymerized polymers serve as scaffolds to encapsulate the cells. An ideal bioink for SLA printing should be biocompatible, non-cytotoxic, and cytocompatible to allow for the attachment and proliferation of the cells. At the same time, bioinks, once crosslinked, should possess sufficient mechanical strengths to retain the 3D-printed structures without collapsing after printing ^[13]^. There are two main categories of photocrosslinkable polymers for bioprinting, namely methacrylated natural polymers and synthetic polymers. Natural polymers, such as gelatin, collagen, hyaluronic acid, and alginate, can be functionalized with methacryloyl groups to enable photopolymerization ^[14–17]^. One of the most commonly-used photocrosslinkable natural polymers is GelMA, a methacrylated form of gelatin. Gelatin is a hydrolyzed form of collagen, which has been shown to support cellular activities as they contain peptide sequences that allow direct binding and remodeling activities from the cells ^[18]^. However, natural polymers tend to have weak mechanical properties ^[19]^, rendering them susceptible to deformation with poor shape fidelity when printed with methacrylated natural polymers alone. On the other hand, synthetic polymers, such as PEG-based polymers, have robust mechanical properties suitable for SLA printing ^[20–21]^. However, they usually do not contain bioactive components to promote cellular attachment or proliferation ^[22]^. In some cases, they may even cause cytotoxicity after printing ^[23–24]^.

Because a single polymer does not fulfill the complete requirement of bioinks, composite inks have been developed using multiple polymers. For example, GelMA and hyaluronic acid methacryloyl (HAMA) were combined to create a composite bioink with increased mechanical stiffness for SLA bioprinting ^[25–26]^. Alternatively, GelMA has been combined with high molecular weight (MW) poly(ethylene glycol) diacrylate (PEGDA) to create composite inks for SLA bioprinting ^[11, 27]^. While low MW PEGDA (MW = 700) is known to compromise the cell viability of the composite ink ^[22, 24]^, high MW PEGDA (MW = 3400) is widely used to formulate SLA-printable bioinks. However, high MW PEGDA is substantially more costly than low MW PEGDA (Supplementary Information; Table S1). In-house synthesis of high MW PEGDA would be possible with specific expertise but may not be readily achievable in every laboratory setting. As such, identifying a material alternative to high MW PEGDA would be beneficial in SLA 3D bioprinting.

### Our approach

To overcome this gap, we aimed to develop an alternative composite bioink for SLA bioprinting that exhibits suitable mechanical properties and cell viability. The developed bioink should compose of materials that are readily accessible (and ideally commercially available). Our previous research revealed that non-cytotoxic PEGDMA (MW = 1000) could be used as an alternative to PEGDA (MW = 700) to create a photocurable ink with porcine GelMA for *in vitro* cell culture ^[24]^. In this work, we explored the suitability of the GelMA-PEGDMA composite ink in SLA 3D bioprinting to create 3D and cell-laden structures with tunable stiffness. We found that adding polyvinyl alcohol (PVA) to the composite bioink enhanced the homogeneity of cell distribution within the 3D-printed constructs. Lastly, the bioprinted construct was shown to support cell proliferation for up to 6 days of culture *in vitro*. Overall, our research identified PEGDMA as a synthetic polymer alternative to high MW PEGDA in SLA bioprinting. The developed composite bioink can be easily formulated, which can potentially be applied for the printing of complex biomimetic structures to create tissue surrogates for therapeutic screening and other biomedical applications.

## Results and Discussion

### Experimental Design

The objective of this work is to develop a bioink that can be used for SLA bioprinting. The overview of this study is summarized (Figure 1). The components of an SLA bioink include live cells, photoinitiators, photoasbsorbers, and polymers. After photocrosslinking, the polymers serve as scaffolds on which the live cells attach and proliferate. To formulate the desirable SLA bioink, the polymer blend should include components that possess suitable mechanical properties to support their own weight during the printing process. At the same time, the formulated SLA bioink should be non-cytotoxic and be permissive to cellular activities, such as cell proliferation.

**Figure 1.**
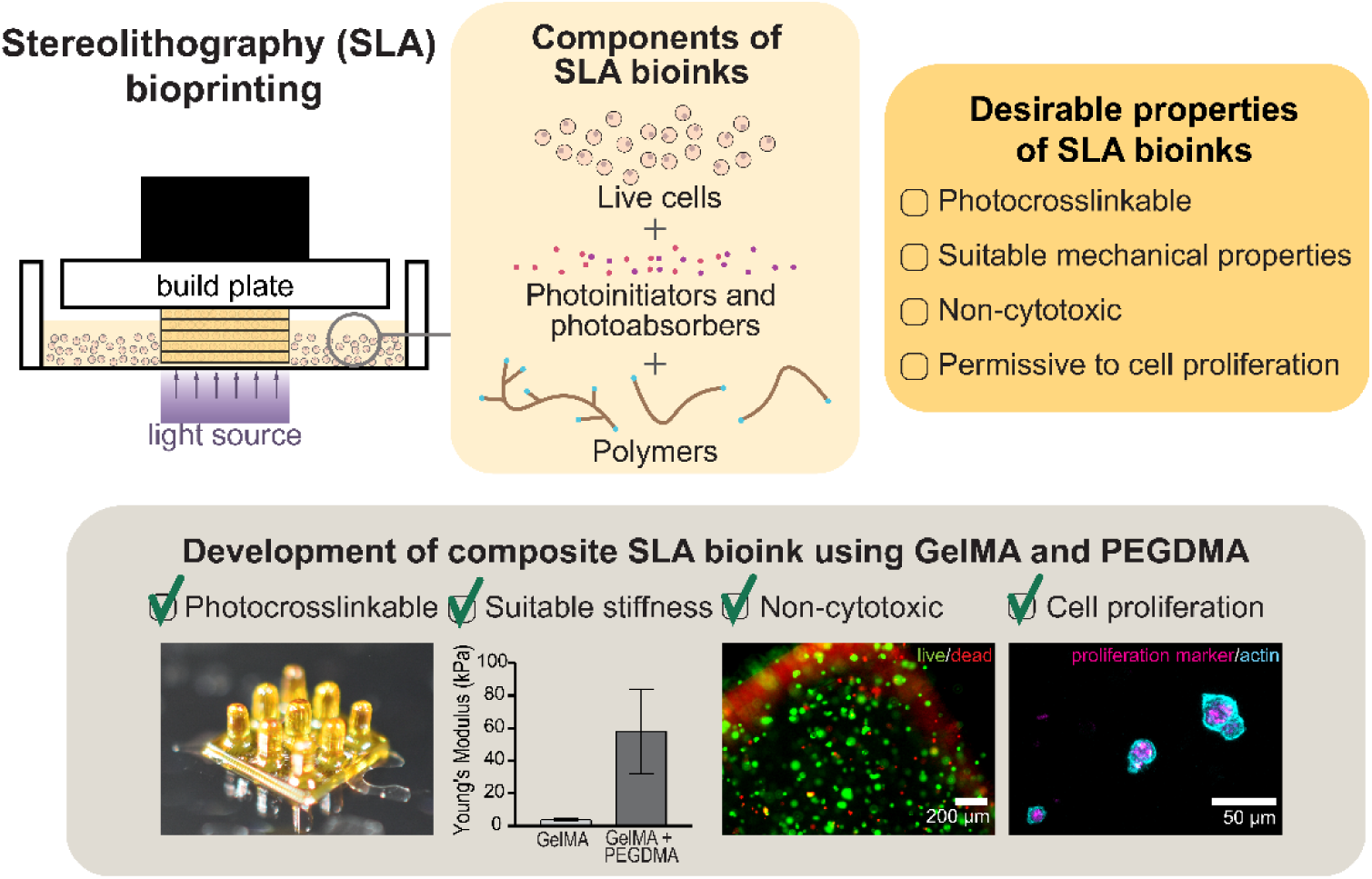
Overview and contribution of this research. The required components in SLA bioink are illustrated. A composite bioink for SLA bioprinting was developed from GelMA and PEGDMA. An ideal bioink should possess tunable mechanical properties upon photocrosslinking and support the viability and proliferation of cells.

In this study, we focused on formulating a composite polymer blend for SLA bioprinitng by combining methacrylated natural polymer and synthetic polymer. We selected cold water fish GelMA as the methacrylated natural polymer. There are multiple sources of gelatin; fish, bovine and porcine are the major sources ^[28]^. However, only fish GelMA remains liquid at room temperature, which is a desirable property for experimental handling during bioprinting. For the synthetic polymer added to the composite ink, we continue to explore the use of PEGDMA 1000 (MW = 1000). Previous research has shown that they are non-cytotoxic and permissive to cellular attachment without peptide modifications ^[29–30]^. PEGDMA contains acrylate moiety that makes them photocurable. These characteristics make PEGDMA 1000 a good candidate for the synthetic additive of the bioinks used in SLA bioprinting.

### Characterization of mechanical properties – 3D-printed GelMA, PEGDMA, and composites

Initially, the mechanical properties of the 3D-printed constructs consisting of relevant materials were characterized. 3D constructs of photocured GelMA (20% w/v) without any additive were mechanically weak with Young’s modulus of 3.85 kPa (Figure 2A). Although GelMA was printable at 20% w/v, shape retention after printing was challenging. Coupled with the high enzymatic degradability of GelMA ^[14, 30]^, constructs consisting of GelMA are known to lose their printed architecture quickly. We also observed that GelMA was not SLA-printable at a concentration below 20% w/v when there were no additives. Here, not-SLA-printable means that no structure appeared on the build plate after printing. Alternatively, constructs printed with 15% v/v PEGDMA exhibited Young’s modulus of 82.8 kPa, which was ∼20-fold higher than the crosslinked GelMA (Figure 2A). Similarly to GelMA, PEGDMA was not printable at a concentration below 15% v/v. Since PEGDMA alone does not allow cellular attachment and proliferation sufficiently ^[29]^, we then explored the printing of the composite inks.

**Figure 2.**
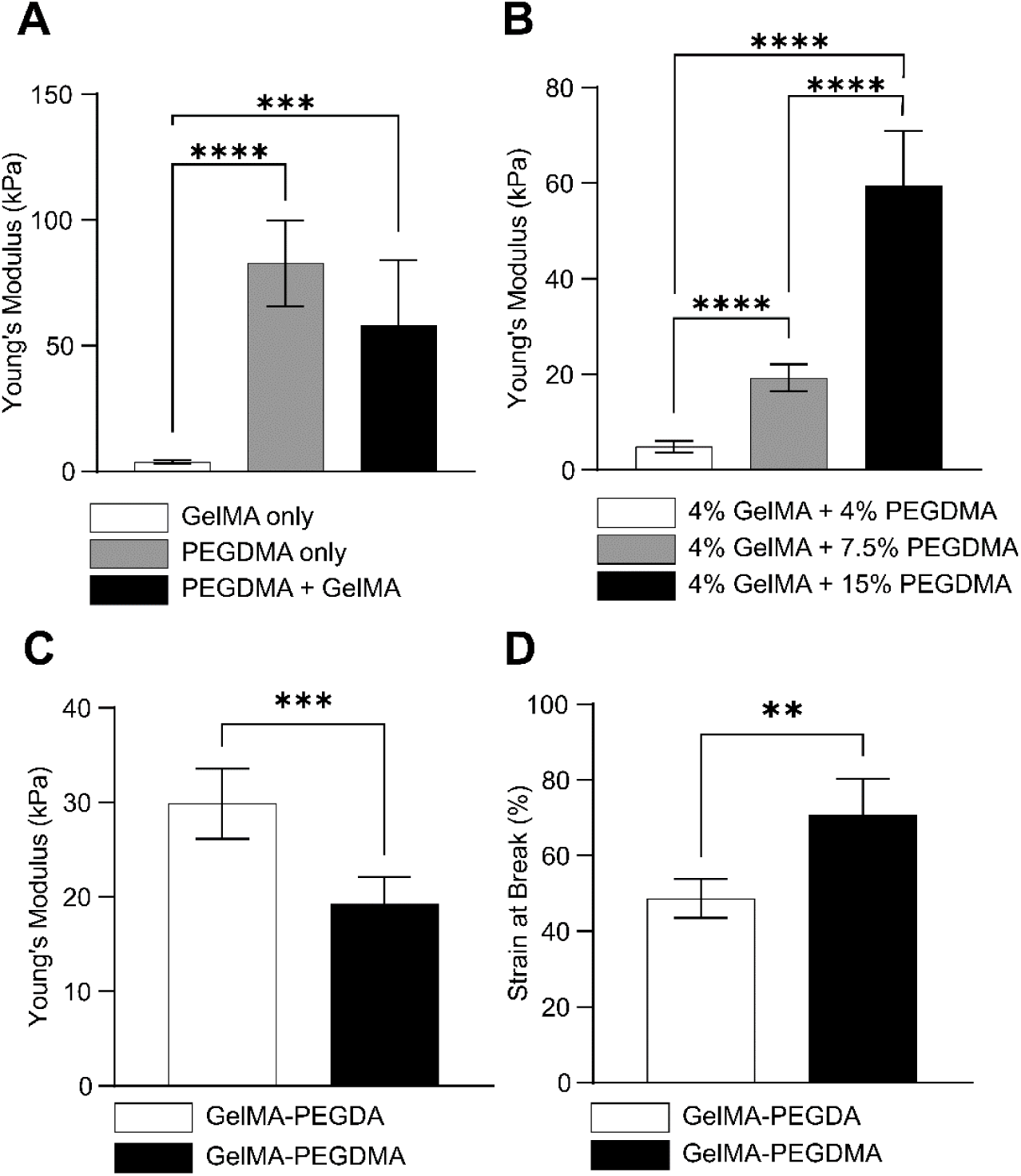
Mechanical characterization of 3D-printed constructs consisting of (1) GelMA, (2) PEGDMA, and (3) GelMA-PEGDMA composite. A) Young’s modulus of cylinders printed using GelMA, PEGDMA, and GelMA-PEGDMA composite inks. B) Young’s modulus of cylinders printed using composite inks with increasing PEGDMA concentrations (*i.e.*, 4% v/v, 7.5% v/v, and 15% v/v). C) Young’s modulus of cylinders printed using GelMA-PEGDA and GelMA-PEGDMA. D) Strain at break of cylinders printed using GelMA-PEGDA and GelMA-PEGDMA. For C) and D), GelMA was at 4% w/v; PEGDA and PEGDMA were at 6% v/v. Data are reported as mean ± SD, n ≥ 5, **p < 0.01, ***p < 0.001, ****p < 0.0001.

The 3D construct consisting of the composite ink of GelMA (6% w/v) and PEGDMA (15% v/v) exhibited intermediate stiffness with Young’s modulus of 58.1 kPa (Figure 2A). Constructs printed using this composite ink displayed good shape retention after printing. By keeping a constant concentration of GelMA at 4% w/v, we found that the PEGDMA concentrations directly correlate to the stiffness of the printed constructs. The increased PEGDMA concentrations resulted in the increased stiffness of the printed constructs (Figure 2B). On the other hand, changing GelMA concentrations in the composite ink (while keeping the PEGDMA concentration as 15% v/v) did not result in significant changes in Young’s modulus of the printed construct (Figure S1). These results suggested that the stiffness of the 3D construct printed by the composite ink was primarily determined by PEGDMA present in ink. Interestingly, when GelMA and PEGDMA were printed as a composite, lower concentrations of both polymers could be used to produce 3D constructs by SLA; for example, GelMA at 4% w/v and PEGDMA at 4% v/v was SLA-printable (Figure 2B), which is far lower than the threshold value to be SLA-printable when a single polymer was used in ink.

We note that PEGDA is a widely used photocurable polymer used to tune the mechanical properties of photocurable inks. Comparing PEGDA (MW = 700) to PEGDMA (MW = 1000) as a supplement in the composite ink containing GelMA (4% w/v), PEGDMA (6% v/v) resulted in less stiff and more elastic constructs than PEGDA (6% v/v) (Figure 2C, 2D). In addition to the non-cytotoxic characteristics of PEGDMA, these results also demonstrated that PEGDMA is a more suitable synthetic photocrosslinkable polymer than PEGDA (MW = 700) as a supplement to GelMA in SLA bioprinting.

### SLA bioprinting using GelMA-PEGDMA composite ink

To demonstrate the printability of GelMA-PEGDMA composite ink using a DLP printer, we printed various structures using the developed ink. Throughout this study, the mixture of PEGDMA (6% v/v) and GelMA (4% w/v) were used; the same formulation was later used for printing with live cells. Figure 3A displays that a distinct sharp peak of a millimeter-scale pyramid was printed. Such sharp peaks was only achievable with high-resolution printing using DLP; the resolution of the pixel of DLP printer used was 80 μm. The same ink was also used to print high-aspect-ratio structures (inspired by villi), where pillars 0.6 mm in height and 0.3 mm in diameter were fabricated. Crucially, we successfully printed overhang structures in the form of a Voronoi, where multiple overhanging pillars 0.5 mm in diameter were fabricated. The capability to print overhangs is important in bioprinting to achieve complex biomimetic architecture such as 3D branching networks found in vasculature. We also investigated the ability to print fine features using the same ink. Figure 3B shows the complex pseudo-2D pattern of the inks (inspired by coral), which demonstrated the printed line width as small as 130 μm. These demonstrations suggested that the GelMA-PEGDMA composite ink could be used to print large millimeter-to-centimeter constructs with 3D features and fine details required to reconstitute physiologically relevant structures.

**Figure 3.**
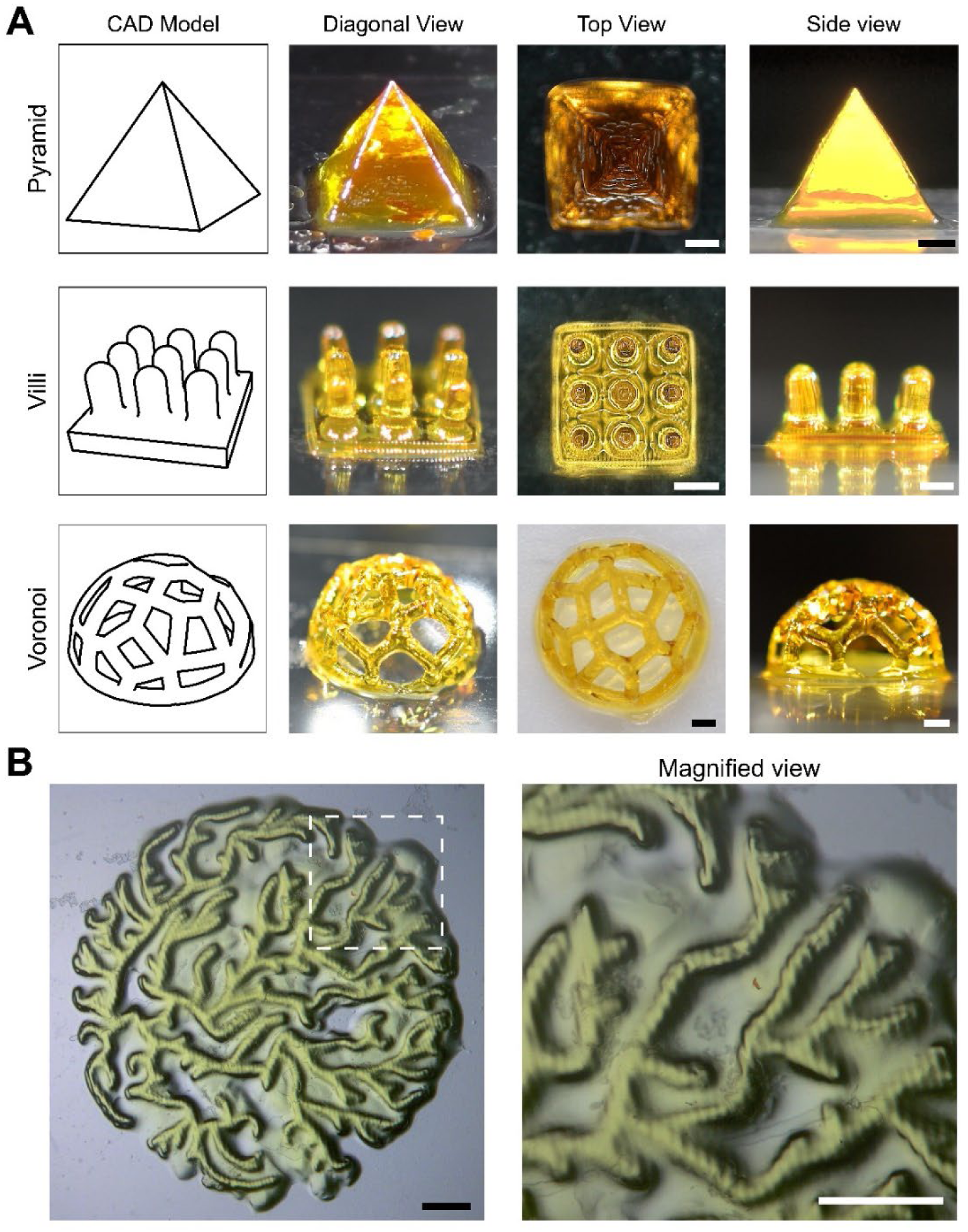
SLA 3D printing of the GelMA-PEGDMA composite ink. A) Pyramid, villi, and Voronoi structures printed using GelMA-PEGDMA composite ink. The first column shows the computer-aided drawing (CAD) model of the structures; the second, third, and fourth columns show the digital photos of each construct taken at orthogonal, top, and side views, respectively. B) Coral structure printed using GelMA-PEGDMA ink. The right image shows a magnified view of the dotted square on the left. Scale bars = 1 mm for A) and B).

### Addition of PVA to enhance homogeneity of cell distribution

In SLA bioprinting, photocrosslinking of the bioink takes place in a layer-by-layer manner starting from the build plate downward; subsequent layers were printed underneath the previously printed layers (Figure 4A). Upon each exposure to light defining each layer of the photocrosslinked structure, the printbed moves upward and then returns downward. Due to this motion of the printbed, low-viscosity bioinks are preferred because the unpolymerized bioink can flow back easily under the lifted build plate for the printing of the subsequent layer ^[31–32]^. However, the low viscosity of the bioink could also lead to sedimentation of cells in the bioink, causing uneven distribution of cells (along the *z*-direction) within the printed construct.

**Figure 4.**
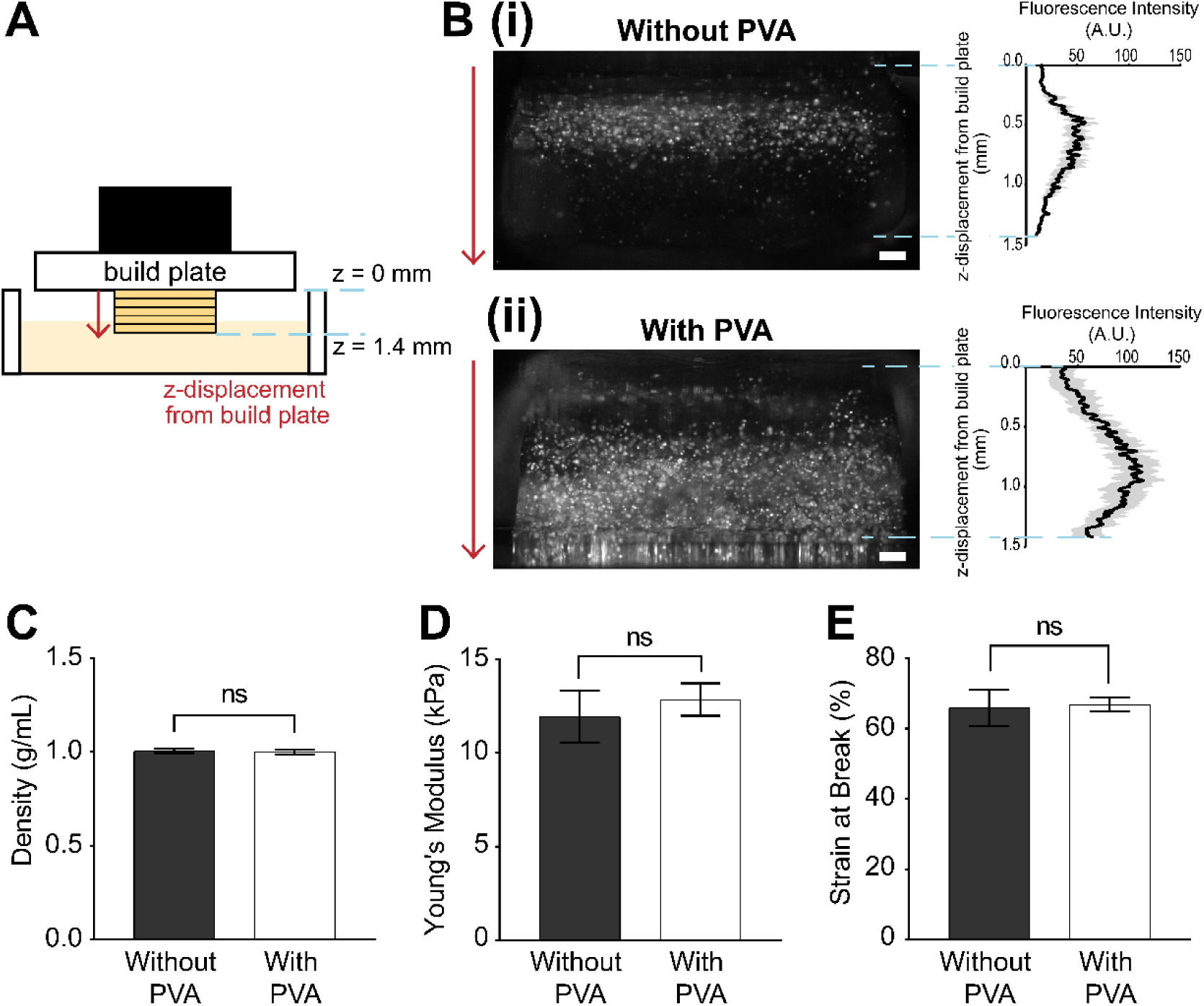
Addition of PVA to increase the homogeneity of the distribution of cells in the GelMA-PEGDMA 3D-printed constructs. A) Illustration showing the concept of SLA printing, where layers of prints are added sequentially to the build plate. B) Fluorescence images of 3D-printed constructs with fluorescent-tagged MDA-MB-231 cells (i) without PVA and (ii) with PVA added to the bioink. The right graphs show the average fluorescence intensity against the corresponding location on the printed block on the left. The location indicated as *z* = 0 cm corresponds to the location of the build plate. C) Density of the bioink with and without PVA. D) Young’s modulus of printed constructs consisting of the bioink with and without PVA. E) Strain at break of printed constructs consisting of the bioink with and without PVA. Data are reported as mean ± SD, n = 5, **p < 0.01, ***p < 0.001, ****p < 0.0001, ns: No statistical significance. Scale bars = 2 mm for B).

To understand the distribution of the printed cells within the constructs, we employed fluorescently labeled cells. First, RFP-tagged MDA-MB-431 cells were embedded in the GelMA-PEGDMA composite ink without additives. The fluorescence intensity profile suggested that the cells were only concentrated in slices up to 1 mm away from the build plate (Figure 4B(i)). In contrast, when PVA (0.1 mg/mL) was added to the bioink to stabilize the cell suspension, the fluorescence intensity profile of the printed construct suggested the presence of the cells up to the slices 1.4 mm away from the build plate (Figure 4B(ii)). This distance (1.4 mm) was the maximum distance that could be captured by the field of view of microscope, and it does not indicate the maximum distance cells can be suspended.

The addition of PVA did not cause a pronounced change in the density of the bioink (Figure 4C). Similarly, the mechanical properties, such as stiffness and elasticity (Figures 4D, 4E) of the printed constructs, were not altered by the addition of PVA at 0.1 mg/mL. When the concentration of PVA was increased, the bioink could not be polymerized with the same printing parameters. PVA absorbes light at 405 nm, albeit at low levels ^[33]^, which is not compatible with the photoinitiator we used in this experiment. The undesired absorbance of light might have caused inefficient photocrosslinking within the bioink and contributed to the failure of SLA bioprinting with increased PVA concentrations. Regardless, our observation suggested that PVA at 0.1 mg/mL was effective in stabilizing cells and achieving homogenous distribution of cells in printed constructs without compromising the mechanical properties of the 3D printed constructs.

### Viability and proliferation of embedded cells

Lastly, we investigated the viability of the embedded cells in the crosslinked composite ink and their capability to proliferate within the crosslinked hydrogel constructs. First, human colorectal cancer cells (SW-480 and Caco-2) were embedded in the GelMA-PEGDMA constructs, and the cell viability was evaluated 2 hr after printing (Day 0), on Day 2 and on Day 6 of cell culture *in vitro* (Figure 5A). It was observed that the cell viability was 79.6% 2 hr after printing (Figure 5B), showing non-cytotoxicity of the composite ink. Although cell viability decreased to 47.9% on Day 2, it did not decrease significantly until Day 6 (Figure 5B). This result shows that the survival of cells stabilized from Day 2 onwards within the printed construct. The decline in cell viability may be attributed to the lack of perfusion within the printed construct. A similar trend of cell viability was also reported in unperfused constructs printed using GelMA-PEGDA composite ink ^[27]^.

**Figure 5.**
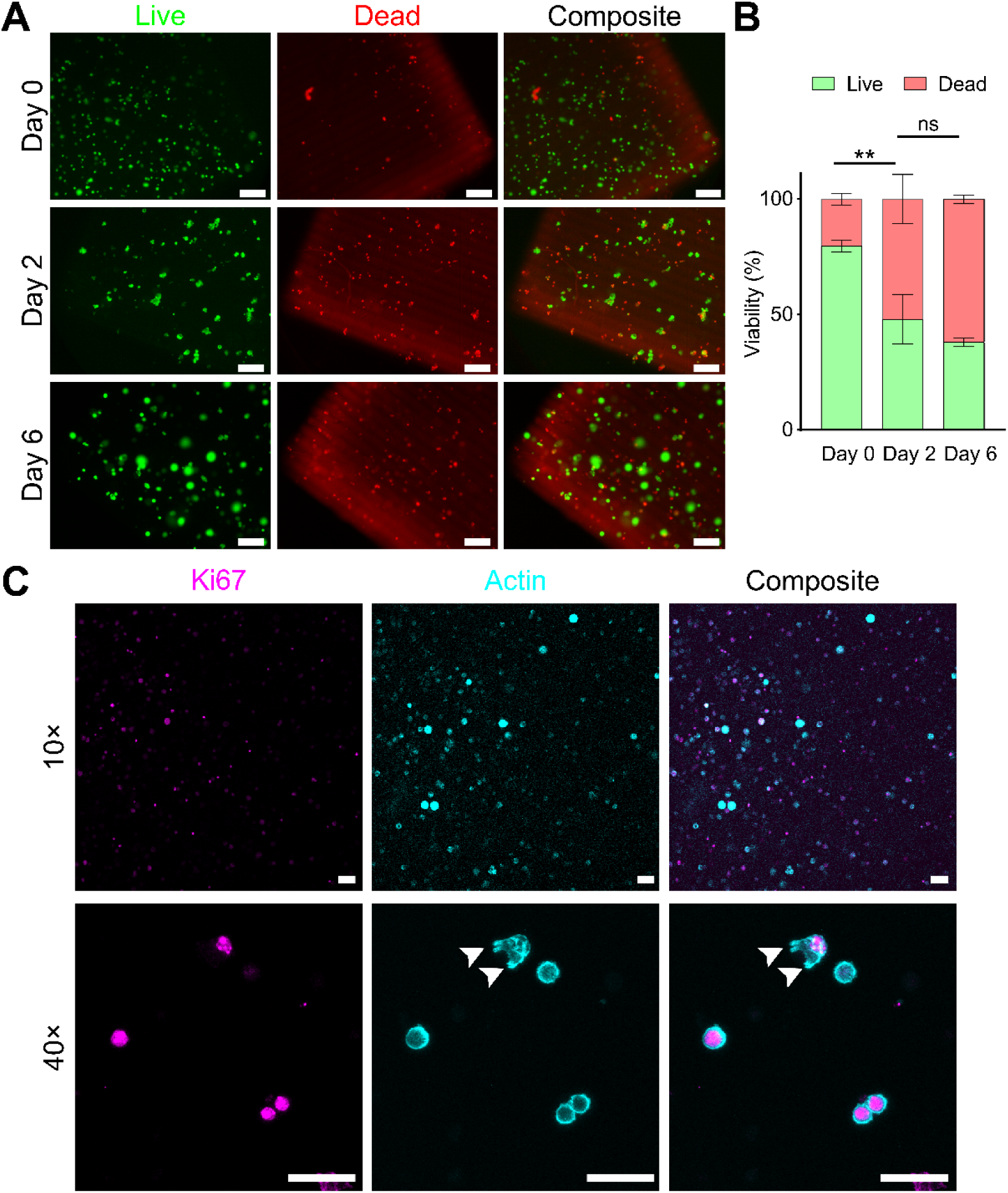
Cell viability and proliferation in the GelMA-PEGDMA composite ink. A) Fluorescence images of live (green) and dead (red) Caco-2 cells in the prints on Days 0, 2, and 6. B) Quantification of live and dead cells on Day 0, 2, and 6. Data are reported as a mean of three fields of view, **p < 0.01. ns: No statistical significance. C) Confocal microscopy images of embedded cells six days after culturing at 10× and 40× magnification. Scale bars = 200 µm for A), 50 µm for C).

We then examined the ability of cells to proliferate within the crosslinked composite ink. The embedded cells appeared as single cells within the printed construct on Day 0, while they appeared in clusters on Day 2 (Figure 5A, Rows 1 and 2). This observation suggested that the embedded cells have proliferated within the construct between Day 0 and Day 2. Cellular staining of Ki67 marker confirmed the presence of mitotic activities within the embedded cells, demonstrating the proliferative capability of embedded cells on Day 6 (Figure 5C). Cellular staining of phalloidin on Day 6 revealed protrusions from the cell (Figure 5C, second row, white arrowhead), suggesting the migration of the cells within the GelMA-PEGDMA construct ^[34]^. Together, these results demonstrated that the GelMA-PEGDMA composite ink can be applied to bioprinting to fabricate large cell-laden constructs with viable and proliferative cells for up to 6 days of culture *in vitro*, which is a substantial culture duration for acute toxicity studies. The cell viability may be enhanced by performing a perfusion culture, which is under current investigation.

## Conclusions

In this research, we validated PEGDMA as a component in bioinks for SLA bioprinting. The addition of PEGDMA to GelMA produced a composite polymer ink suitable for bioprinting using a DLP printer to produce 3D constructs with tunable stiffness (Young’s modulus of ∼5 – 60 kPa). The GelMA-PEGDMA composite ink could be used to produce printed structures with intricate geometry and architecture (including overhang features), which demonstrated the ability to mimic the complex structure of the biological constructs. Adding PVA in the bioink enhanced the homogeneity of the cell distribution within the printed construct along the direction of the printing (i.e., *z*-direction). Importantly, the GelMA-PEGDMA composite ink was biocompatible with low cytotocicity and sustained the proliferative ability of cells embedded in the bioprinted constructs. With simple and low-cost formulation, the developed GelMA-PEGDMA ink will make bioprinting more accessible to researchers with diverse expertise and accelerate the progress of tissue engineering research. The high-resolution and flexible printing enabled by low-cost DLP printers combined with our readily available bioinks will eventually enable biofabrication of complex cell-laden biomimetic structures, such as the villi structures in the intestines, or interconnecting vasculature, for tissue engineering and the study of regenerative medicine. The ability to encapsulate viable cancer cells within the 3D-printed constructs should also bring out the potential of our bioinks in applications for patient-specific cancer models for therapeutic screening.

## Materials and Methods

### Materials

PEGDA 700 (average molecular weight, MW = 700) and tartrazine were purchased from Sigma-Aldrich (St. Louis, MO, USA). PEGDMA 1000 (average molecular weight, MW = 1000) was purchased from Polysciences (Washington, PA, USA). Lyophilized fish GelMA was purchased from Gelomics (Brisbane, QLD, Australia). Ruthenium (Ru) and sodium persulfate (SPS) were purchased from Advanced Biomatrix (San Diego, CA, USA). Poly(vinyl alcohol) (PVA) Grade 1788L (MW = 72600 − 81400, degree of hydrolysis = 85% − 90%) was purchased from Eastchem (Qingdao, China).

### Bioink preparation

All PEG-based polymers (PEGDA 700 and PEGDMA 1000) were prepared in volume-per-volume (v/v) accordingly in phosphate buffered saline (PBS) (Nacalai Tesque, Kyoto, Japan) to obtain solutions of their respective concentrations. 100% v/v PEGDMA 1000 was warmed to 37 °C to obtain a liquid form to prepare PEGDMA stock solution (80% v/v) with PBS for the rest of the experiments. Fish GelMA was dissolved in PBS at 37 °C under constant stirring on a heat plate to prepare 10% w/v stock solution, and 20% w/v solution for printing with GelMA alone, with PBS for each experiment. The fish GelMA stock solution was stored and used within one week. RU and SPS were used as photoinitiators at a final concentration of 0.2 and 2 mM, respectively, while tartrazine was used as photoabsorber at 1.2 mM. In all bioprinting with cells, cell density in the bioink was 6 million cells per millimeter.

### *3D* printing procedure

All 3D printing were performed on Photon Ultra (Anycubic, Shenzhen, China) with 25 μm layer height with an exposure time of 15 s per layer; the pixel resolution of the printer was 80 μm. The light source provided light at a wavelength of 405 nm with an intensity of 4.5 mW/cm^2^. All prints were printed on coverslips attached to the build plate of the printer. The printed constructs were detached from coverslips using a alcohol-sterilized microtome blade. All prints were designed in Rhinoceros (Robert McNeel & Associates, WA, USA) and sliced using Photon Workshop (Anycubic, Shenzhen, China).

### Measurement of mechanical properties of the bioinks

To characterize the compressive modulus and deformability of the 3D-printed constructs using different polymer blends, cylinders of 4 mm in diameter and 2 mm in height were printed using each polymer blend for compression tests. Compression tests were carried out using the Dynamic Mechanical Analyzer TA Q800 (TA instruments, New Castle, DE, USA). Tests were carried out with a preload force of 0.001 N to ensure conformity between the top plate and surface of the sample. A ramp force rate of 0.4 N min^-1^ up to 10 N was applied at room temperature. The compressive modulus was determined as the slope of the linear region corresponding with 0 – 20% strain.

### Cell culture maintenance

Cell lines used in this work were red fluorescence protein-tagged (RFP) human breast adenocarcinoma cell line, MDA-MB-231 (GenTarget Inc, San Diegom, CA, USA), RFP-tagged human colorectal adenocarcinoma cell line, SW480 (Angio-Proteomie, Boston, MA, USA), and Caco-2 human colorectal adenocarcinoma cell line (ATCC, Manassas, VA, USA). All cell lines were maintained in Dulbecco’s Modified Eagle (DMEM) high glucose (Nacalai Tesque, Kyoto, Japan) supplemented with fetal bovine serum (at 10% v/v for MDA-MB-231, SW480, and at 20% v/v for Caco-2), 1× Antibiotic-Antimycotic (Gibco, Waltham, USA) and 0.1% v/v MycoZap (Lonza, Basel, Switzerland). Media change was performed every other day. Cells were incubated in 37 °C incubator with 5% carbon dioxide. Dissociation of cells from the tissue culture flasks (Greiner, Kremsmünster, Austria) was performed by first rinsing the cells twice with PBS before incubating the cells in trypsin (Nacalai Tesque, Kyoto, Japan) for 3 min in 37 °C incubator. Trypsinized cells were spun down at 1000 rpm for 3 min before getting neutralized using DMEM growth media.

#### Cell viability evaluation

Square blocks (2.7 mm × 2.7 mm × 0.7 mm) of Caco-2 cells were printed using the print parameters stated in the previous sections. The printed constructs were submerged in cell culture media and on a sea-saw shaker rocking at 2 rpm and incubated for maintenance in 37 °C incubator with 5% carbon dioxide. The cell-laden constructs were rinsed once with PBS before incubating in live/dead assay reagents (Invitrogen, Waltham, MA, USA) for 30 min at 37 °C. The cell-laden constructs were then imaged using a fluorescence microscope (Zeiss Axio Observer D1, Oberkochen, Germany) and quantified using ImageJ ^[35]^.

### Immunostaining of cell-laden printed constructs

The printed constructs were rinsed thrice using PBS for 10 min per wash. They were then fixed using 4% w/v paraformaldehyde (PFA) in PBS for 40 min on a see-saw shaker at 2 rpm. Permeabilization was performed using 1% Triton-X-100 in PBS over two nights at 4 °C and blocked over one night using blocking buffer (1% Triton-X-100, 2% bovine serum albumin (BSA), and 0.2% NaN_3_ in PBS) at 4 °C. Anti-Ki-67 antibody (ab15580, Abcam, Cambridge, UK) was used at 1:50 dulition with Alexa Fluor^TM^ 488 Phalloidin (A12379, Invitrogen, Waltham, MA, USA) at 1:100 ratio in antibody dilution buffer (0.2% Triton-X-100, 2% BSA, and 0.2% NaN_3_ in PBS) over 2 nights at 4 °C. Washings was performed thrice with 1 hr per wash at room temperature for the first two washes, the last wash was overnight at 4 °C on a see-saw rocker. Next, Donkey anti-rabbit secondary antibody (A31573, Invitrogen, Waltham, MA, USA) at 1:500 dilution and SYTOX green (S7020, Invitrogen, Waltham, MA, USA) at 1:10000 dilution in antibody dilution buffer were used for incubation over one night. Secondary antibody washings were performed twice at 120 min per wash at room temperature before one last wash overnight at 4 °C on a see-saw rocker.

The samples were then washed thrice with PBS before being submerged in RapiClear (Hsinchu City, Taiwan) overnight. The samples were then imaged on a confocal microscope (LSM700, ZEISS, Oberkochen, Germany).

## Acknowledgements

M.H. acknowledges Agency for Science, Technology and Research (A*STAR) (A*STAR-AMED joint grant, A19B9b0067) and MOE, Singapore (Academic Research Fund (AcRF) Tier 2, MOE2019-T2-2-192) for the project funding.

## Declarations

- The authors declare compliance with ethical standards.
- The authors declare that there is no conflict of interest.
- The authors declare that this study does not contain any studies with human or animal subjects performed by any of the authors.

## Notes

### Competing Interest Statement

The authors have declared no competing interest.

## References

[1] S. V. Murphy, A. Atala, Nat Biotechnol 2014, 32, 773.

[2] S. V. Murphy, P. De Coppi, A. Atala, Nat Biomed Eng 2020, 4, 370.

[3] Y. S. Zhang, G. Haghiashtiani, T. Hübscher, D. J. Kelly, J. M. Lee, M. Lutolf, M. C. McAlpine, W. Y. Yeong, M. Zenobi-Wong, J. Malda, *Nat. Rev.* Methods Primers 2021, 1.

[4] A. Blaeser, D. F. Duarte Campos, U. Puster, W. Richtering, M. M. Stevens, H. Fischer, Adv Healthc Mater 2016, 5, 326.

[5] S. Boularaoui, G. Al Hussein, K. A. Khan, N. Christoforou, C. Stefanini, Bioprinting 2020, 20.

[6] H. Cui, M. Nowicki, J. P. Fisher, L. G. Zhang, Adv Healthc Mater 2017, 6.

[7] G. Burke, D. M. Devine, I. Major, Polymers 2020, 12.

[8] T. Ching, J. Vasudevan, S. Y. Chang, H. Y. Tan, A. Sargur Ranganath, C. T. Lim, J. G. Fernandez, J. J. Ng, Y. C. Toh, M. Hashimoto, *Small* 2022, 18, e2203426.

[9] D. Gonzalez-Hernandez, S. Varapnickas, A. Bertoncini, C. Liberale, M. Malinauskas, Adv. Opt. Mater. 2022, 11.

[10] K. Parkatzidis, M. Chatzinikolaidou, M. Kaliva, A. Bakopoulou, M. Farsari, M. Vamvakaki, ACS Biomater Sci Eng 2019, 5, 6161.

[11] B. Grigoryan, S. J. Paulsen, D. C. Corbett, D. W. Sazer, C. L. Fortin, A. J. Zaita, P. T. Greenfield, N. J. Calafat, J. P. Gounley, A. H. Ta, F. Johansson, A. Randles, J. E. Rosenkrantz, J. D. Louis-Rosenberg, P. A. Galie, K. R. Stevens, J. S. Miller, Science 2019, 364, 458.

[12] R. Zhang, N. B. Larsen, Lab Chip 2017, 17, 4273.

[13] J. Gopinathan, I. Noh, Biomater. Res. 2018, 22.

[14] M. Zhu, Y. Wang, G. Ferracci, J. Zheng, N. J. Cho, B. H. Lee, Sci Rep 2019, 9, 6863.

[15] N. Pien, D. Pezzoli, J. Van Hoorick, F. Copes, M. Vansteenland, M. Albu, B. De Meulenaer, D. Mantovani, S. Van Vlierberghe, P. Dubruel, Mater Sci Eng C Mater Biol Appl 2021, 130, 112460.

[16] B. S. Spearman, N. K. Agrawal, A. Rubiano, C. S. Simmons, S. Mobini, C. E. Schmidt, J Biomed Mater Res A 2020, 108, 279.

[17] M. Hasany, S. Talebian, S. Sadat, N. Ranjbar, M. Mehrali, G. G. Wallace, M. Mehrali, Appl. Mater. Today 2021, 24.

[18] K. Yue, G. Trujillo-de Santiago, M. M. Alvarez, A. Tamayol, N. Annabi, A. Khademhosseini, Biomaterials 2015, 73, 254.

[19] M. C. Catoira, L. Fusaro, D. Di Francesco, M. Ramella, F. Boccafoschi, J Mater Sci Mater Med 2019, 30, 115.

[20] P. Joshi, S. Breaux, J. Naro, Y. Wang, M. S. U. Ahmed, K. Vig, M. L. Auad, J. Appl. Polym. Sci. 2021, 138.

[21] J. P. Mazzoccoli, D. L. Feke, H. Baskaran, P. N. Pintauro, J Biomed Mater Res A 2010, 93, 558.

[22] V. Chan, P. Zorlutuna, J. H. Jeong, H. Kong, R. Bashir, Lab Chip 2010, 10, 2062.

[23] G. Liu, Y. Li, L. Yang, Y. Wei, X. Wang, Z. Wang, L. Tao, RSC Adv. 2017, 7, 18252.

[24] S.-Y. Chang, T. Ching, M. Hashimoto, Mater. Today Proc. 2022, 70, 179.

[25] R. Hossain Rakin, H. Kumar, A. Rajeev, G. Natale, F. Menard, I. T. S. Li, K. Kim, Biofabrication 2021, 13.

[26] M. Wang, W. Li, J. Hao, A. Gonzales, 3rd, Z. Zhao, R. S. Flores, X. Kuang, X. Mu, T. Ching, G. Tang, Z. Luo, C. E. Garciamendez-Mijares, J. K. Sahoo, M. F. Wells, G. Niu, P. Agrawal, A. Quinones-Hinojosa, K. Eggan, Y. S. Zhang, Nat Commun 2022, 13, 3317.

[27] N. Anandakrishnan, H. Ye, Z. Guo, Z. Chen, K. I. Mentkowski, J. K. Lang, N. Rajabian, S. T. Andreadis, Z. Ma, J. A. Spernyak, J. F. Lovell, D. Wang, J. Xia, C. Zhou, R. Zhao, Adv Healthc Mater 2021, 10, e2002103.

[28] J. Alipal, N. A. S. Mohd Pu’ad, T. C. Lee, N. H. M. Nayan, N. Sahari, H. Basri, M. I. Idris, H. Z. Abdullah, Mater. Today Proc. 2021, 42, 240.

[29] R. R. Asawa, J. C. Belkowski, D. A. Schmitt, E. M. Hernandez, A. E. Babcock, C. K. Lochner, H. N. Baca, C. M. Rylatt, I. S. Steffes, J. J. VanSteenburg, K. E. Diaz, D. M. Doroski, PLoS One 2018, 13, e0202825.

[30] C. B. Hutson, J. W. Nichol, H. Aubin, H. Bae, S. Yamanlar, S. Al-Haque, S. T. Koshy, A. Khademhosseini, Tissue Eng Part A 2011, 17, 1713.

[31] S. Bertlein, G. Brown, K. S. Lim, T. Jungst, T. Boeck, T. Blunk, J. Tessmar, G. J. Hooper, T. B. F. Woodfield, J. Groll, Adv Mater 2017, 29.

[32] A. Schwab, R. Levato, M. D’Este, S. Piluso, D. Eglin, J. Malda, Chem Rev 2020, 120, 11028.

[33] H. Abral, M. Ikhsan, D. Rahmadiawan, D. Handayani, N. Sandrawati, E. Sugiarti, A. N. Muslimin, J. Mater. Res. Technol. 2022, 17, 2193.

[34] P. T. Caswell, T. Zech, Trends Cell Biol. 2018, 28, 823.

[35] J. Schindelin, I. Arganda-Carreras, E. Frise, V. Kaynig, M. Longair, T. Pietzsch, S. Preibisch, C. Rueden, S. Saalfeld, B. Schmid, J. Y. Tinevez, D. J. White, V. Hartenstein, K. Eliceiri, P. Tomancak, A. Cardona, Nat Methods 2012, 9, 676.

