## Supplementary Information for "3D Bioprinting Using Poly(ethylene-glycol)-dimethacrylate (PEGDMA) Composite"

Restricted

Restricted

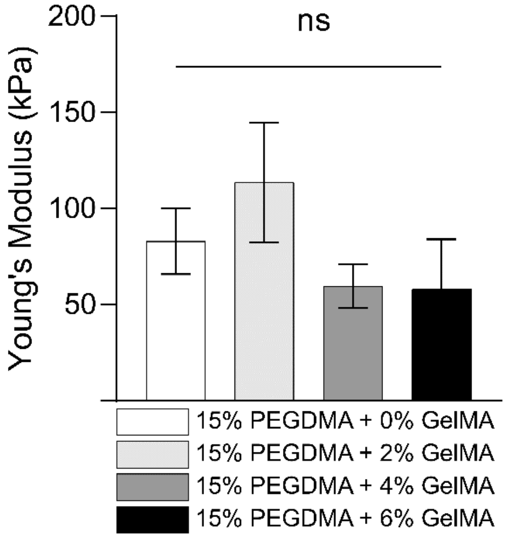


**Figure S1.** Young's modulus of printed constructs using various composite inks with GelMA concentrations (0 – 6% w/v). The concentration of PEGDMA was fixed at 15% v/v. ns: No statistical difference

Restricted

**Table S1.** Price comparison of PEGDA and PEGDMA of various MW.

| Synthetic Polymer | Molecular Weight (g) | Cost per gram (USD) | Source |
|---|---|---|---|
| **Poly(ethylene glycol) diacrylate (PEGDA)** | 575 | 0.52 | Sigma |
| | 700 | 0.50 | Sigma |
| | 1000 | 210.00 | Advanced Biomatrix |
| | 3400 | 80.00 | Allevi |
| | | 210.00 | Advanced Biomatrix |
| | 6000 | 178.00 | Sigma |
| | | 80.00 | Allevi |
| | | 210.00 | Advanced Biomatrix |
| **Poly(ethylene glycol) dimethacrylate (PEGDMA)** | 750 | 0.32 | Sigma |
| | 1000 | 85.40 | Sigma |
| | | 4.90 | Polyscience |
| | 4000 | 285.00 | Sigma |
| | 6000 | 397.00 | Sigma |
| | 8000 | 55.00 | Polyscience |

Restricted
